# Contrasting Mitochondrial Diversity of Endemic *Corbicula* Clams in Sulawesi’s Ancient Lakes: Phylogeography and Implications for Conservation

**DOI:** 10.64898/2026.07.02.735996

**Authors:** Gunawan Muhammad, Bayu K. A. Sumarto, Diky Dwiyanto, I Gede Jose Dewana, Andi Chadijah, Septiana Sri Astuti, Asep Sahidin, Thomas von Rintelen

## Abstract

The global study of freshwater clams in the genus *Corbicula* is frequently confounded by invasive androgenetic lineages that experience mitochondrial DNA capture and clonal propagation. In contrast, the endemic *Corbicula* of Sulawesi’s ancient lakes reproduce sexually, offering a uniquely reliable system for mitochondrial population genetics. This study provides the first population-genetic framework for two endemic species, *Corbicula possoensis* (Lake Poso) and *C. linduensis* (Lake Lindu), using the cytochrome *c* oxidase subunit I (COI) marker. We analysed 90 newly generated COI sequences from *C. possoensis* (six stations) and *C. linduensis* (three stations), integrated with reference sequences from GenBank, to assess genetic diversity, population structure, and phylogeographic patterns. Hierarchical AMOVA revealed deep divergence between the two lakes (Φ_CT = 0.607), consistent with prolonged independent isolation rather than a single shared vicariance event, as the two species do not form a sister pair in the phylogeny. Within Lake Poso, *C. possoensis* exhibited exceptionally high genetic diversity (24 haplotypes; *h* = 0.876; π = 0.016) and pronounced micro-geographic structuring into three phylogeographic zones (North: Tentena and Siuri; East: Tando Nceppo and Busogo Beach; Southwest: Bancea and Pendolo), each characterised by distinct haplogroups. Remarkably, the maximum divergence between zones (K2P = 2.33%) approached the interspecific distance between *C. possoensis* and *C. linduensis* (K2P = 2.42%), indicating that within-lake mitochondrial divergence has reached near-interspecific levels. Conversely, *C. linduensis* displayed near-panmixia and extreme genetic depauperation (3 haplotypes; *h* = 0.246; π = 0.0004), indicating long-term demographic stasis within a restricted habitat. The deep phylogeographic zonation in *C. possoensis* suggests that its discrete populations should be treated as separate Management Units (MUs) in conservation planning to preserve locally adapted gene complexes, whereas the severely depauperate gene pool of *C. linduensis* renders it critically vulnerable to environmental disturbance and invasive species, warranting urgent IUCN Red List assessment. To validate these mitochondrial boundaries and inform future conservation strategies, multi-marker and genome-wide reassessments are strongly recommended.

## 1. Introduction

Freshwater bivalves represent one of the most imperilled faunal groups on Earth, with extinction rates far exceeding those of any comparable taxon (Lydeard et al. 2004, Lopes-Lima et al. 2021). Despite their ecological role as ecosystem engineers and bioindicators, conservation efforts for these organisms are chronically hampered by fundamental gaps in population-level genetic data — information critical for delineating management units, identifying refugia, and guiding translocation programmes (Frankham 2010, Lopes-Lima et al. 2017). Within this broader crisis, the genus *Corbicula* (Megerle von Mühlfeld, 1811) occupies a uniquely paradoxical position in freshwater malacology. Globally, the genus has attracted considerable molecular attention, yet this body of work has been dominated by the study of invasive lineages — particularly *Corbicula fluminea* (O. F. Müller, 1774) and allied taxa — whose reproductive biology fundamentally undermines the utility of standard mitochondrial markers (Pigneur et al. 2011, Kropotin et al. 2023). These invasive clams reproduce via androgenesis, a form of clonal inheritance in which the maternal nuclear genome is discarded and offspring develop from paternal genetic material alone (Komaru et al. 1998, Hedtke et al. 2008, Pigneur et al. 2012). A key consequence of androgenesis is ‘mitochondrial capture’, wherein paternal mitochondrial genomes can be transmitted across deeply divergent nuclear backgrounds (Pigneur et al. 2014). A related phenomenon can also arise through introgressive hybridisation independent of androgenesis, further complicating mitochondrial genealogies (Pfenninger et al. 2002). The result is that COI haplotype networks for androgenetic *Corbicula* specifically reflect the genealogy of mitochondrial genomes rather than organismal phylogeny, rendering population structure inferences from COI alone unreliable or misleading in these lineages (Park and Kim 2003, Vastrade et al. 2022). This pervasive mitochondrial unreliability has long cast a shadow over population genetic studies of the entire genus. However, this issue is not universal. It is largely confined to studies of invasive lineages such as *C. fluminea* and *C. leana*, in which androgenesis has been confirmed even in native Southeast Asian populations through the presence of biflagellate spermatozoa (Pootanon et al. 2026). The pervasive focus on these androgenetic taxa has obscured the existence of reproductively distinct lineages within the same genus that are amenable to standard mitochondrial analysis.

A critical but underappreciated exception to this pattern exists among the endemic *Corbicula* fauna of the ancient lakes of Sulawesi, Indonesia. Lakes Poso and Lindu, together with the Malili Lake system, harbour a unique radiation of lacustrine *Corbicula* that diverges profoundly from their invasive congeners in reproductive biology. Anatomical and ultrastructural studies have confirmed that *Corbicula possoensis* (Sarasin & Sarasin, 1898) and *Corbicula linduensis* (Billinger, 1914), along with other Sulawesian lacustrine species, possess monoflagellate spermatozoa and maintain a dioecious, sexually reproducing reproductive mode (Glaubrecht et al. 2003, Korniushin and Glaubrecht 2003). This fundamental biological distinction carries a profound methodological implication: because androgenesis and the associated phenomenon of mitochondrial capture are absent, standard mitochondrial DNA markers behave as expected under conventional population genetic theory in these taxa. Sequence divergence reflects genealogical history, haplotype structure reflects population connectivity, and diversity indices carry interpretable biological meaning. Accordingly, the ancient lake endemics of Sulawesi represent a substantially more tractable system for mitochondrial population genetics within this otherwise problematic genus. Although introgressive hybridisation cannot be entirely excluded in the absence of nuclear data, the elimination of androgenesis removes the primary source of mitochondrial–nuclear discordance documented in invasive lineages — making these taxa a uniquely informative, if not unconditionally reliable, system that has paradoxically remained entirely unexploited for population-level analysis.

Despite their scientific value and conservation urgency, both *C. possoensis* and *C. linduensis* remain among the most genetically data-deficient endemic bivalves in Wallacea. The only molecular study of these taxa (von Rintelen and Glaubrecht 2006) employed several COI sequences per species as representative terminals in a genus-level phylogeny — a sampling design sufficient to establish interspecific relationships but wholly inadequate for inferring intraspecific diversity, population structure, or demographic history. The data deficit is especially acute for *C. linduensis*: the sole entry attributed to this species in GenBank (acc. no. DQ285579, deposited as part of Glaubrecht, 2006) has since been reidentified as *C. leana* based on a molecular-phylogenetic analysis (Bespalaya et al. 2025), leaving the species without a single valid, publicly accessible COI sequence. This taxonomic ambiguity is not isolated: a recent integrative reassessment using COI barcoding and morphology across a broad geographic range has further clarified species boundaries within the genus, reassigning several Southeast Asian sequences previously attributed to endemic taxa to the invasive *C. leana* lineage (Bespalaya et al. 2026). Taken together, these revisions underscore both the instability of historical identifications and the urgent need for fresh population-level sampling of the true lacustrine endemics.

The molecular invisibility is compounded by unresolved taxonomic standing — MolluscaBase currently subsumes *C. linduensis* under *Corbicula moltkiana* var. *lindoensis*, inhibiting its recognition as an independent conservation unit — and by its categorisation as a ‘missing’ species by the The Recently Extinct Plants and Animals Database (2024). Meanwhile, *C. possoensis* has been assessed as Endangered (EN) on the IUCN Red List (Rintelen and Bogan 2012), reflecting a restricted range and documented habitat degradation within Lake Poso (Sulawesty et al. 2024). For both species, the complete absence of population genetic baselines represents a critical obstacle to any evidence-based conservation strategy.

Here, we present the first population genetic assessment of *C. possoensis* and *C. linduensis* based on mitochondrial COI sequences, explicitly leveraging the sexual reproductive mode of these taxa to justify the reliability of this marker for intraspecific inference. Specifically, this study aims to: (i) characterise haplotype diversity and nucleotide diversity within and among sampling stations across Lake Poso (*C. possoensis*, six stations, n = 60) and Lake Lindu (*C. linduensis*, three stations, n = 30); (ii) assess population structure and inter-station connectivity using haplotype networks, analysis of molecular variance (AMOVA), and pairwise FST statistics; and (iii) evaluate demographic history through neutrality tests to identify signatures of population expansion or contraction. Together, these data constitute the first genetic baseline for both species and provide empirical foundations for conservation prioritisation across the two lake systems.

## 2. Materials and Methods

### 2.1. Ethical permits

All field activities were conducted under institutional ethical approval from the National Research and Innovation Agency (BRIN) ethics commission, in compliance with Indonesian national regulations. Approvals were granted for Animal Care and Use (No. 130/KE.02/SK/07/2025) and for the Handling of Hazardous Biological Materials (No. 59/KE.06/SK/08/2025). Research endorsement was additionally obtained from the Scientific Authority and Biodiversity Secretariat (SKIKH) of BRIN (No. B-2785/IV/KS.00/4/2025). For field activities in Lake Lindu, which falls under active customary governance, authorisation was secured from the *Tua Adat* (customary elder) of the *To‘oma To‘Lindu* indigenous community.

### 2.2. Sample collections

Specimens of *C. possoensis* were collected from six stations distributed along the littoral zone of Lake Poso: Tentena (P6; 1°46.4’S, 120°38.3’E), Tando Nceppo (P2; 1°53.2’S, 120°39.6’E), Siuri (P14; 1°48.3’S, 120°31.7’E), Busogo Beach (P4; 1°59.7’S, 120°41.1’E), Bancea (P11; 1°58.6’S, 120°35.1’E), and Pendolo (P9; 2°03.9’S, 120°41.2’E). Specimens of *C. linduensis* were collected from three stations along the littoral zone of Lake Lindu: Nangkating (L1; 1°20.9’S, 120°04.8’E), Uwe Rawa (L2; 1°16.9’S, 120°06.2’E), and Pulau (L3; 1°20.8’S, 120°04.1’E). Sampling in Lake Lindu was limited to three of six planned stations owing to *Ombo*, a customary prohibition enforced by the *To’oma To’Lindu* community; access to the three sampled stations was granted through permission from the *Tua Adat* (customary elder). Sampling locations are shown in Fig. 1.

**Fig. 1.**
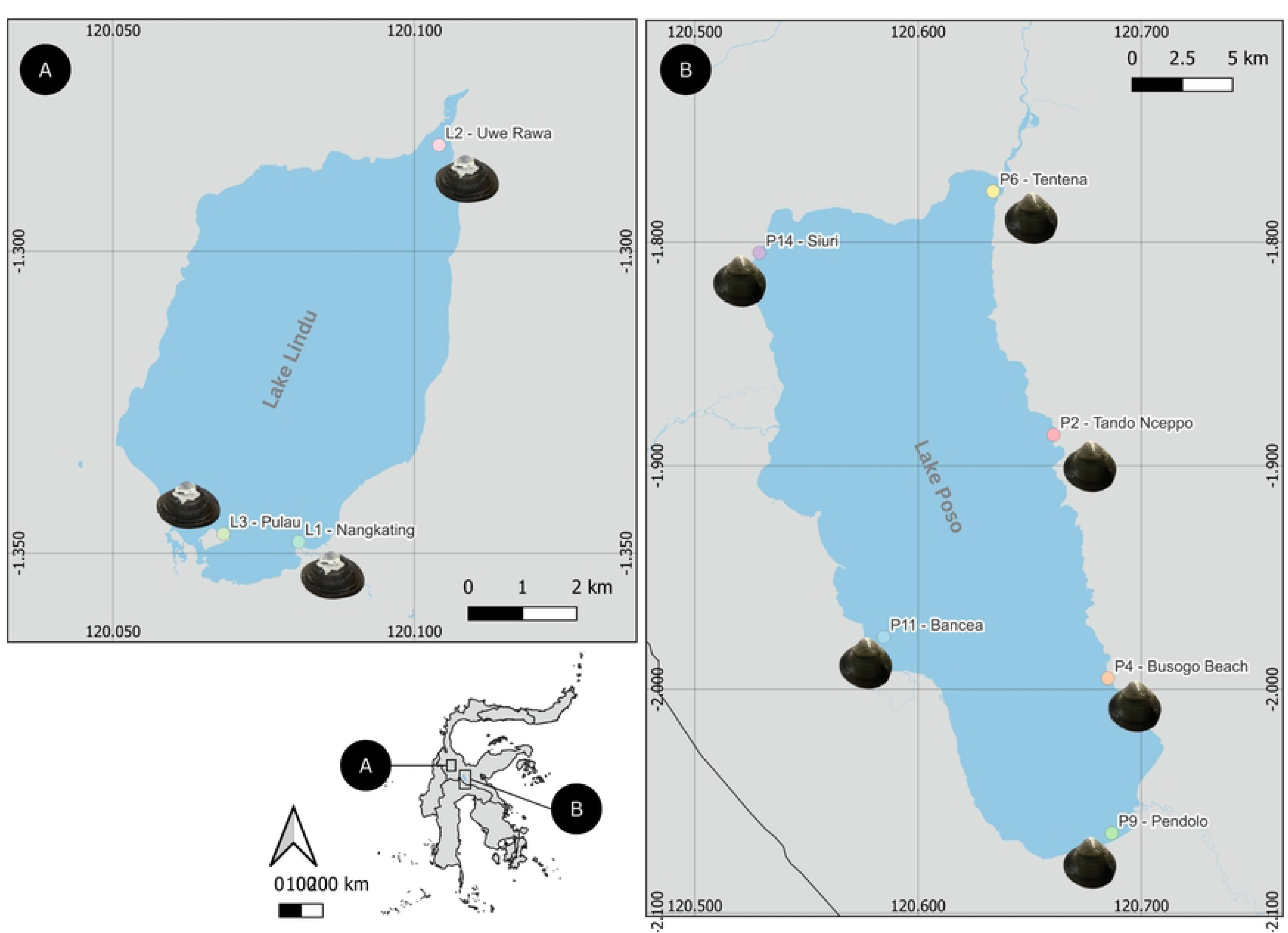
Sampling localities for *Corbicula possoensis* and *Corbicula linduensis*. A): Lake Lindu, showing the three sampling stations of *C. linduensis*: Nangkating (L1), Uwe Rawa (L2), and Pulau (L3). B): Lake Poso, showing the six sampling stations of *C. possoensis*: Tentena (P6), Tando Nceppo (P2), Siuri (P14), Busogo Beach (P4), Bancea (P11), and Pendolo (P9).

At each station, ten adult specimens per species were collected by 15 x 15 cm Ekman grab and hand-picking along the littoral substrate at depths of 2–15 m during low-activity periods. Foot tissue (ca. 2 mm^3^) was dissected immediately in the field and preserved in 96% ethanol pending molecular analysis. Voucher specimens (shells with preserved soft tissue) were retained for morphological reference and deposited at the Museum Zoologicum Bogoriense (MZB) and Hydrobiology Laboratory, National Research and Innovation Agency (BRIN), Indonesia.

### 2.3. DNA extraction, amplification, and sequencing

Total genomic DNA was extracted from ethanol-preserved foot tissue using the Geneaid DNA Mini Kit (GT300; Geneaid Biotech, New Taipei City, Taiwan) following the manufacturer’s tissue extraction protocol. Extracted DNA was quantified spectrophotometrically and stored at −20°C until further use.

A 710-bp fragment of the mitochondrial Cytochrome c Oxidase subunit I (COI) gene was amplified by polymerase chain reaction (PCR) using the freshwater mussel-optimised primer pair LCO22me2 (5’-GGTCAACAAATCATAAAGATATTGG-3’) and HCO700dy2 (5’-GGRGGRTASACGTTCAGGGTG-3’) (Walker et al. 2006). PCR reactions (35 µL total volume) contained 17.5 µL of 2× Dream Taq PCR Master Mix (Thermo Scientific), 0.5 µL of each primer (10 µM), 2 µL of template DNA (80–120 ng/µL, as quantified by NanoDrop spectrophotometry), and 14.5 µL of nuclease-free water. Thermal cycling conditions were: initial denaturation at 95°C for 5 min; 37 cycles of denaturation at 94°C for 30 s, annealing at 52°C for 30 s, and extension at 72°C for 45 s; with a final extension at 72°C for 5 min. PCR products were visualised on 2% agarose gels stained with ethidium bromide.

Bidirectional Sanger sequencing of purified PCR products was performed by First Base Laboratories (Sdn. Bhd., Selangor, Malaysia). Forward and reverse chromatograms were inspected, trimmed, and assembled into consensus sequences using Geneious Prime. All sequences are deposited in the NCBI GenBank database (Supplementary Table 1).

### 2.4. Sequence alignment and dataset composition

Newly generated COI sequences from this study comprised 60 sequences of *Corbicula possoensis* (six stations, Lake Poso), 30 sequences of *Corbicula linduensis* (three stations, Lake Lindu;), three sequences of *Corbicula matannensis* (Lake Matano;), and three sequences of *Corbicula anomioides* (Dulumai, Lake Poso). The latter two were included to provide phylogenetic context for the population-level analyses.

These sequences were combined with 46 homologous COI sequences retrieved from NCBI GenBank, selected to represent the broader taxonomic and geographic diversity of the genus *Corbicula*. The NCBI dataset comprised two groups. First, representatives of all other known Sulawesian endemic taxa: *C. possoensis* (DQ285597–DQ285599, Lake Poso), *C. anomioides* (DQ285604–DQ285605, Lake Poso), *C. matannensis* (DQ285593–DQ285595, DQ285584, DQ285586, DQ285589, and AY275664; Malili Lake

System), *C. loehensis* (DQ285580–DQ285581, OM912055, Malili Lake System), and sequences previously attributed to *C. subplanata* (DQ285601–DQ285603, Sulawesi rivers; von Rintelen & Glaubrecht, 2006) but subsequently reidentified as *C. leana* (Bespalaya et al. 2025). An additional sequence deposited as *C. linduensis* (DQ285579, Palu River drainage; von Rintelen & Glaubrecht, 2006), also reidentified as *C. leana* (Bespalaya et al. 2025), was included to assess its phylogenetic position relative to true lacustrine *C. linduensis*.

Second, non-Sulawesian reference taxa were included to contextualise the phylogenetic position of the Sulawesian endemic clade: *C. tobae* (MN746824–MN746826, Lake Toba; Kropotin et al. 2022), *C. fluminea* (six sequences from Panama, Malaysia, USA, Russia, Myanmar, and Japan), *C. fluminalis* (three sequences from India), *C. leana* (two sequences from France, one from Japan), *C. javanica* (two sequences from Indonesia), *C. sandai* (AF196273), *C. blandiana* (PV023642, Thailand), *C. elatior* (OM698379, Russia), *C. moltkiana* (three sequences from Indonesia), *C. largillierti* (PV023648, Myanmar), *C. lamarckiana* (PV023665, Laos), and *C. africana* (OM912292, South Africa). *Corbicula japonica* (AB845593) was designated as the outgroup, following Pootanon et al. (2026). Multiple sequence alignment was performed using the MUSCLE algorithm implemented in Geneious Prime. The final alignment comprised 142 sequences of 590 bp. A complete list of accession numbers and specimen metadata is provided in Supplementary Table S1.

### 2.5. Phylogenetic analyses

The best-fit substitution model for the COI alignment was selected using ModelFinder (Kalyaanamoorthy et al. 2017) as implemented in IQ-TREE 3 v3.1.2 (Minh et al. 2020), evaluating candidate DNA models. HKY+F+G4 was selected as the best-fit model under the Bayesian Information Criterion (BIC; BIC score = 5,247.19), while TN+F+I+G4 was preferred under AIC criteria. The BIC-selected model, HKY+F+G4, was applied for maximum likelihood (ML) tree inference, as BIC is recommended for datasets of moderate size where parameter parsimony is prioritised. ML inference was conducted with 10,000 ultrafast bootstrap replicates (UFBoot; Hoang et al. 2018) and 10,000 replicates of the SH-aLRT branch test. Nodes receiving UFBoot ≥ 95% and SH-aLRT ≥ 80% were considered well-supported.

### 2.6. Haplotype network

Median-joining haplotype networks were constructed using Hapsolutely (Vences et al. 2024) at two complementary levels: (i) separate intraspecific networks for *C. possoensis* and *C. linduensis* to maximise resolution of within-species haplotype relationships, and (ii) a combined network incorporating all available Sulawesian *Corbicula* taxa to assess interspecific haplotype affinities and evaluate the phylogenetic placement of the focal species within the regional radiation. Haplotypes were colour-coded by sampling station in the intraspecific networks, and by taxon in the combined network, to facilitate visual assessment of geographic and taxonomic structure.

### 2.7. Genetic diversity indices

For each species, the following genetic diversity statistics were calculated per station and across all stations combined using custom R scripts executed in RStudio, with the ape (Paradis and Schliep 2019) and pegas (Paradis 2010) packages: number of haplotypes (*H*), haplotype diversity (*h*), nucleotide diversity (π), number of polymorphic sites (*S*), and average number of pairwise nucleotide differences (*k*). Between-zone pairwise genetic distances were calculated as both uncorrected p-distances and distances under the Kimura two-parameter model (K2P; Kimura, 1980) using the ape package in R (model = ‘raw’ and model = ‘K80’, respectively; Paradis and Schliep, 2019). p-distance is reported as the primary conservative measure; K2P values are provided for comparison.

### 2.8. Population genetic structure

Hierarchical analysis of molecular variance (AMOVA) was performed using the pegas package (Paradis 2010) in R v.4.4.0 to partition total genetic variance into components attributable to: (i) differences between lakes (*C. possoensis* vs *C. linduensis*), (ii) differences among stations within each lake, and (iii) differences among individuals within stations. Statistical significance was assessed using 10,000 permutations. Pairwise genetic differentiation between all station pairs within each lake was estimated as Φ∼ST∼ (Weir and Cockerham 1984) using the same framework, with significance assessed by 10,000 permutations and Bonferroni correction for multiple comparisons (α = 0.0033 for 15 pairs in *C. possoensis*; α = 0.0167 for 3 pairs in *C. linduensis*).

### 2.9. Isolation by distance

The relationship between genetic differentiation [*F*_ST_/(1 − *F*_ST_)] and the natural logarithm of geographic distance (km) between station pairs was tested separately for each lake using a Mantel test (Mantel 1967, Rousset 1997) implemented in the vegan package (Oksanen et al. 2022) in R v.4.4.0, with 9,999 permutations. Geographic distances between stations were calculated from GPS coordinates using the Haversine formula.

### 2.10. Demographic history

Signatures of historical population expansion or contraction were assessed for each species using: (i) Tajima’s *D* statistic (Tajima 1989), calculated via tajima.test() in the pegas package (Paradis, 2010) with significance assessed under the normal approximation; and (ii) Fu’s *F*∼S∼ statistic (Fu 1997), calculated using a custom implementation of the Ewens sampling formula with significance assessed via the right-tail probability *S* (significant if *S* < 0.02). Both statistics were also computed per sampling station to characterise within-lake spatial heterogeneity in demographic signals. Mismatch distributions of pairwise sequence differences were computed using a custom R function and compared against the expected distribution under a sudden population expansion model (Rogers and Harpending 1992); model fit was assessed using the raggedness index *r* (Harpending 1994) and the sum of squared deviations (SSD). All demographic analyses used the ape (Paradis and Schliep 2019) and pegas packages in R v.4.4.0.

## 3. Results

### 3.1. Sequence characteristics and dataset composition

The final COI alignment comprised 142 sequences of 590 bp, representing 96 newly generated sequences from this study and 46 sequences retrieved from NCBI GenBank. Mean gap and ambiguous character content across all sequences was 0.01%, and no sequence failed the chi-squared nucleotide composition test (*p* > 0.05 for all 142 sequences), confirming compositional homogeneity across the dataset. Of the 590 alignment positions, 422 were constant (71.5%), 55 were parsimony-uninformative singletons (9.3%), and 113 were parsimony-informative (19.2%), yielding 120 distinct site-pattern combinations.

### 3.2. Phylogenetic relationships

The ML phylogenetic tree (Fig. 2), rooted on *C. japonica* (AB845593), resolves three well-supported subclades within the Lake Poso assemblage and recovers *C. linduensis* as a clearly distinct, monophyletic lineage (SH-aLRT/UFBoot = 89.2/98). The Lake Poso clade — encompassing *C. possoensis* sequences from all six stations alongside *C. anomioides* — receives modest support at its root node (SH-aLRT/UFBoot = 82.1/75), directly reflecting the non-monophyletic arrangement of *C. possoensis* across the tree. Three internal *C. possoensis* subclades are recovered: Subclade I (SH-aLRT/UFBoot = 98.8/99), predominantly comprising Siuri (P14) and Tentena (P6) specimens; Subclade II (SH-aLRT/UFBoot = 84.1/99), comprising Bancea (P11) and Pendolo (P9) specimens; and Subclade III (SH-aLRT/UFBoot = 85/99), comprising peripheral haplotypes from Busogo Beach (P4) and Tando Nceppo (P2). A fourth group of sequences from P2, P9, P11, and P6 received insufficient bootstrap support (SH-aLRT/UFBoot = 73.5/66) and remained unresolved, interspersed with *C. anomioides* sequences. The three *C. anomioides* specimens from Dulumai form a well-supported monophyletic subclade (SH-aLRT/UFBoot = 96.5/98) nested within this aforementioned unresolved portion alongside *C. possoensis* sequences. Sequences previously attributed to *C. subplanata* (DQ285601–DQ285603) and *C. linduensis* (DQ285579) cluster within the *C. leana* clade, well-separated from all Sulawesian lacustrin sequences.

**Fig. 2.**
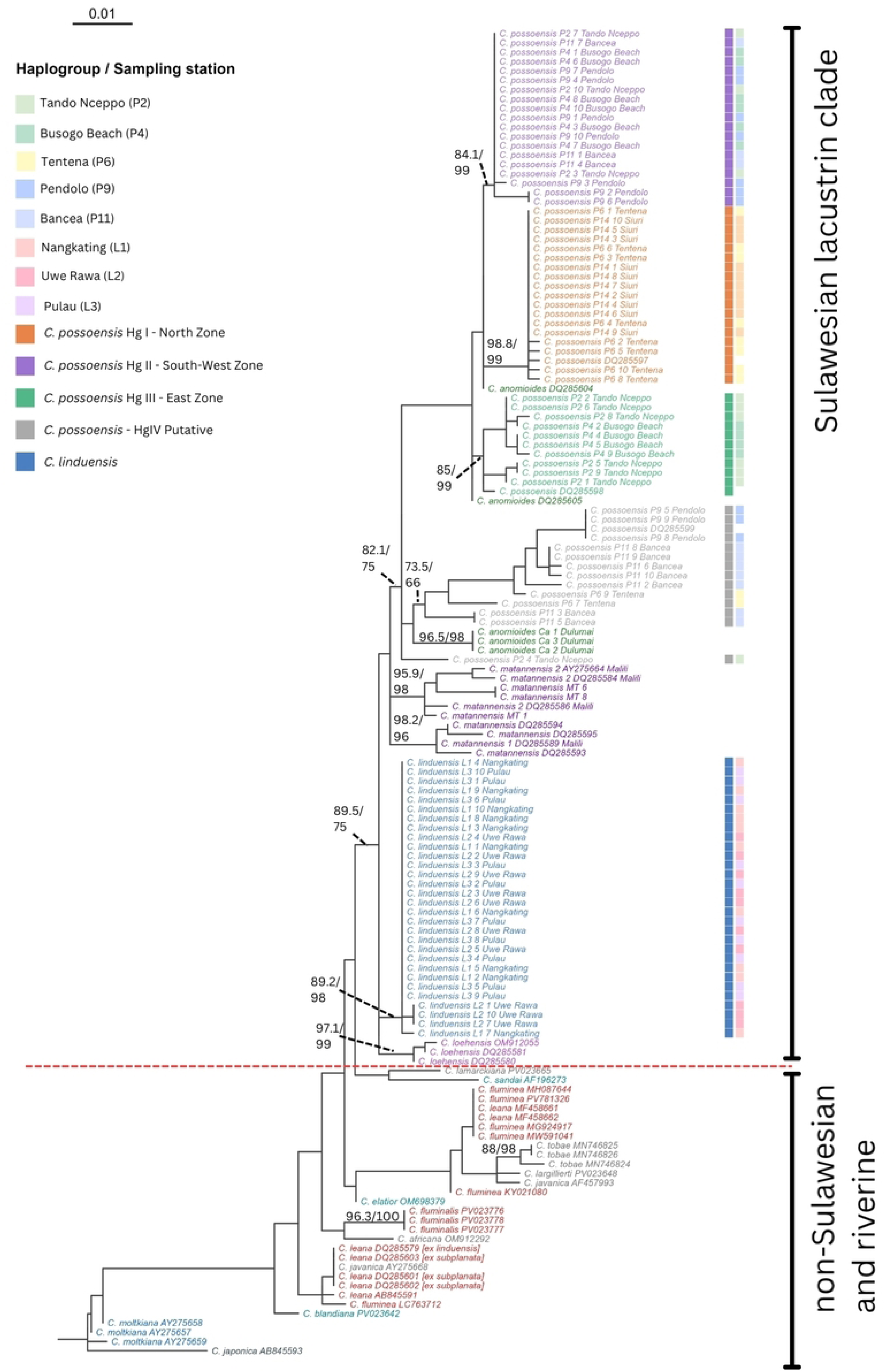
Maximum likelihood (ML) phylogenetic tree inferred from COI sequences (590 bp; *n* = 142) of endemic Sulawesian *Corbicula* and related taxa, reconstructed under the HKY+F+G4 substitution model (BIC = 5,247.19) in IQ-TREE 3 v3.1.2, rooted on *Corbicula japonica* (AB845593). Numbers at internal nodes represent SH-aLRT (%)/UFBoot (%) support values; nodes with SH-aLRT ≥ 80% and UFBoot ≥ 95% are considered well-supported. Coloured bars adjacent to tip labels indicate haplogroup assignment (left bar; *C. possoensis* and *C. linduensis* only) and sampling station (right bar). Haplogroup boundaries were defined based on the median-joining haplotype network (Fig. 3). Reference sequences retrieved from GenBank are indicated by their accession numbers; sequences previously attributed to *C. subplanata* or *C. linduensis* but subsequently reidentified as *C. leana* (Bespalaya et al. 2025) are denoted with brackets. Scale bar represents 0.01 substitutions per site.

### 3.3. Haplotype networks

Haplotype network analysis of *C. linduensis* (*n* = 30) identified three haplotypes (N*h* = 3), with a simple star-like topology and uniformly low diversity (*h* = 0.246; π = 0.000429; Table 1; Fig. 3A). A single dominant haplotype (*n* = 26) was shared across all three sampling stations — Nangkating (L1), Uwe Rawa (L2), and Pulau (L3) — and accounted for 86.7% of all sampled individuals. Specimens from Pulau (L3) were completely monomorphic, with all ten individuals carrying the dominant haplotype. Two minor haplotypes were detected: one exclusive to Uwe Rawa (L2; *n* = 3) and one singleton from Nangkating (L1; *n* = 1), each separated from the central haplotype by a single mutational step. The shallow network topology and low nucleotide diversity of *C. linduensis* are fully consistent with the strongly supported monophyletic clade recovered in the ML phylogeny (SH-aLRT = 89.2%; UFBoot = 98%; Section 3.2), indicating a genetically cohesive population with minimal geographic structuring at the COI locus.

**Fig. 3.**
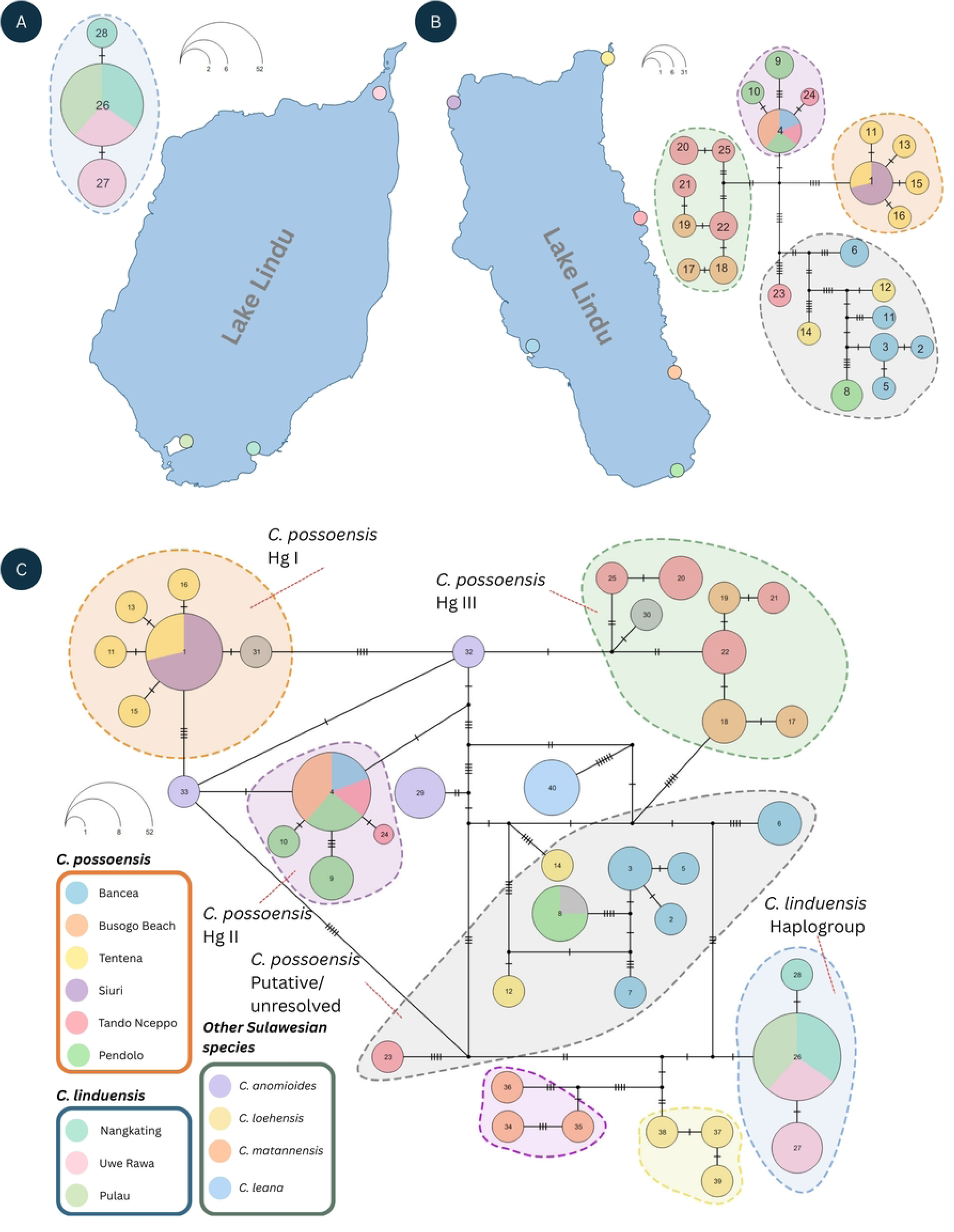
Median-joining haplotype networks of *Corbicula* COI sequences (590 bp) constructed in Hapsolutely. Numbers within circles indicate haplotype identifiers; circle size is proportional to haplotype frequency (see scale indicators); tick marks on connecting lines indicate the number of mutational steps between haplotypes (each tick = 1 step); small, filled circles represent median vectors (inferred ancestral or unsampled haplotypes). Dashed ellipses delineate haplogroup boundaries. (A) Haplotype network of *C. linduensis* (*n* = 30; *Nh* = 3) across three sampling stations in Lake Lindu (Nangkating L1, Uwe Rawa L2, Pulau L3), dominated by a single haplotype (H26; *n* = 26) shared across all stations. (B) Haplotype network of *C. possoensis* (*n* = 60; *Nh* = 24) across six sampling stations in Lake Poso, showing four haplogroups (Hg I–IV) colour-coded by sampling station. (C) Combined median-joining network of all Sulawesian *Corbicula* COI sequences (*n* = 108), including *C. possoensis* (Hg I–IV and putative/unresolved haplotypes), *C. linduensis*, *C. anomioides*, *C. matannensis*, *C. loehensis*, and sequences previously attributed to *C. subplanata* but reidentified as *C. leana* (Bespalaya et al. 2025). Notably, *C. anomioides* occupies a central position in the network, connecting *C. possoensis* haplogroups I, II, and III, consistent with recent divergence between these two sympatric species. Haplotype numbers in panels (B) and (C) are consistent; sampling station colours follow the legend at left.

**Table 1.** Haplotype and nucleotide diversity indices for *Corbicula possoensis* (Lake Poso) and *C. linduensis* (Lake Lindu) per sampling station. *n*, number of sequences; N*h*, number of haplotypes; *h*, haplotype diversity; π, nucleotide diversity; *k*, mean number of pairwise differences; *S*, number of segregating sites.

| Species | Stn. | Locality | Lake | n | Nh | h | $\pi$ | k | S |
| --- | --- | --- | --- | --- | --- | --- | --- | --- | --- |
| *C. possoensis* | P2 | Tando Nceppo | Poso | 10 | 6 | 0.888<br>9 | 0.010094 | 5.956 | 19 |
|  | P4 | Busogo Beach | Poso | 10 | 4 | 0.644<br>4 | 0.005989 | 3.533 | 8 |
|  | P6 | Tentena | Poso | 10 | 7 | 0.866<br>7 | 0.012580 | 7.422 | 27 |
|  | P9 | Pendolo | Poso | 10 | 4 | 0.777<br>8 | 0.016949 | 10 | 22 |
|  | P11 | Bancea | Poso | 10 | 6 | 0.888<br>9 | 0.018041 | 10.64<br>4 | 25 |
|  | P14 | Siuri | Poso | 10 | 1 | 0 | 0.000000 | 0 | 0 |
|  | — | Lake Poso | Poso | 60 | 24 | 0.875<br>7 | 0.016084 | 9.489 | 52 |
| *C. linduensis* | L1 | Nangkating | Lindu | 10 | 2 | 0.2 | 0.000339 | 0.2 | 1 |
|  | L2 | Uwe Rawa | Lindu | 10 | 2 | 0.466<br>7 | 0.000791 | 0.467 | 1 |
|  | L3 | Pulau | Lindu | 10 | 1 | 0 | 0.000000 | 0 | 0 |
|  | — | Lake Lindu | Lindu | 30 | 3 | 0.246 | 0.000429 | 0.253 | 2 |

In stark contrast, the haplotype network of *C. possoensis* (*n* = 60) was markedly more complex, comprising 24 haplotypes (N*h* = 24) with high haplotype and nucleotide diversity (*h* = 0.8757; π = 0.016084; Table 1; Fig. 3B). Four haplogroups were delimited based on network topology, corresponding to the three well-supported subclades and one unresolved peripheral group identified in the ML phylogeny (Section 3.2). Haplogroup I (Hg I; *n* = 19) was anchored by the most frequent haplotype in the dataset (*n* = 14), shared among all ten specimens from Siuri (P14) and four specimens from Tentena (P6), with four satellite haplotypes connected by single mutational steps. Haplogroup II (Hg II; *n* = 19) was centred on a second high-frequency haplotype (*n* = 16) drawn from stations distributed across the eastern (Busogo Beach [P4] and Tando Nceppo [P2]), southern (Pendolo [P9]), and western (Bancea [P11]) shores. Haplogroup III (Hg III; *n* = 11) comprised seven peripheral haplotypes exclusive to Busogo Beach (P4) and Tando Nceppo (P2), consistent with the geographically restricted subclade (SH-aLRT = 85%; UFBoot = 99%) in the ML tree. The putative/unresolved haplogroup (Hg IV; *n* = 14) comprised 9 haplotypes distributed without geographic cohesion across Tentena (P6), Bancea (P11), Pendolo (P9), and Tando Nceppo (P2). Notably, Busogo Beach (P4) and Tando Nceppo (P2) each harboured specimens from multiple haplogroups (Hg II and Hg III, with Hg IV additionally present at P2), indicating pronounced intra-population haplogroup heterogeneity at these localities. Siuri (P14) was the only station exhibiting complete haplogroup monomorphism, with all ten individuals belonging to Hg I.

The combined Sulawesian haplotype network (Fig. 3C) placed *C. possoensis* and *C. linduensis* in two entirely separate, non-overlapping clusters with no shared haplotypes or connecting median vectors, confirming their genetic distinctness at the COI marker. The reticulate, high-diversity cluster of *C. possoensis* contrasted sharply with the compact, low-diversity *C. linduensis* cluster.

### 3.4. Population genetic structure

Hierarchical AMOVA of the combined COI dataset (*n* = 90; six Lake Poso stations and three Lake Lindu stations) revealed that the overwhelming majority of total genetic variance was explained by differences between the two lake systems (Φ_CT_ = 0.607; 60.7% of variance; *p* < 0.001). An additional 16.7% of variance was attributable to differentiation among stations within each lake (Φ_SC_ = 0.423; *p* < 0.001), while only 22.7% of variance resided within stations (Table 2). Overall differentiation across all hierarchical levels was very high (Φ_ST_ = 0.773; *p* < 0.001), consistent with deep genetic discontinuity between *C. possoensis* and *C. linduensis* and pronounced within-lake structuring.

**Table 2.** Pairwise ΦST values (upper triangle) and corresponding *p*-values from permutation tests (lower triangle; 10,000 replicates) for (A) *C. possoensis* [P2 (Tando Nceppo), P4 (Busogo Beach), P6 (Tentena), P9 (Pendolo), P11 (Bancea), P14 (Siuri); six stations, Lake Poso; Bonferroni α = 0.0033 for 15 comparisons] and (B) *C. linduensis* [L1 (Nangkating), L2 (Uwe Rawa), L3 (Pulau); three stations, Lake Lindu; Bonferroni α = 0.0167 for 3 comparisons]. Asterisks (*) indicate pairs significant after Bonferroni correction. Analyses performed using hierarchical AMOVA (Excoffier et al., 1992).

| *(A) Corbicula possoensis — Lake Poso* |  |  |  |  |  |  |
| --- | --- | --- | --- | --- | --- | --- |
|  | P2 | P4 | P6 | P9 | P11 | P14 |
| P2 | — | -0.094 | 0.213 | 0.260 | 0.528* | 0.615* |
| P4 | 0.948 | — | 0.196 | 0.254 | 0.554* | 0.783* |
| P6 | 0.011 | 0.019 | — | 0.168 | 0.501 | 0.043 |
| P9 | 0.016 | 0.023 | 0.073 | — | 0.085 | 0.462* |
| P11 | 0.001 | 0.001 | 0.005 | 0.179 | — | 0.746* |
| P14 | 0.000 | 0.000 | 0.483 | 0.000 | 0.000 | — |
| *(B) Corbicula linduensis — Lake Lindu* |  |  |  |  |  |  |
|  | L1 | L2 | L3 |  |  |  |
| L1 | — | 0.275 | -<br>0.000 |  |  |  |
| L2 | 0.126 | — | 0.222 |  |  |  |
| L3 | 1.000 | 0.208 | — |  |  |  |

Within Lake Poso, *C. possoensis* exhibited significant and substantial inter-station differentiation (Φ_ST_ = 0.404; 40.4% of variance among stations; *p* < 0.001). Pairwise Φ_ST_ values ranged from −0.094 to 0.783 across the 15 station pairs (Table 2). After Bonferroni correction for multiple comparisons (α = 0.0033 for 15 pairs), six pairs were statistically significant: Siuri (P14) vs. Bancea (P11; Φ_ST_ = 0.746), Busogo Beach (P4; Φ_ST_ = 0.783), Tando Nceppo (P2; Φ_ST_ = 0.615), and Pendolo (P9; Φ_ST_ = 0.462), and additionally Bancea (P11) vs. Tando Nceppo (P2; Φ_ST_ = 0.528) and Busogo Beach (P4; Φ_ST_ = 0.554). Conversely, the Siuri–Tentena pair (P14–P6; Φ_ST_ = 0.043; *p* = 0.483) and the Bancea–Pendolo pair (P11–P9; Φ_ST_ = 0.085; *p* = 0.179) showed no significant differentiation, concordant with both pairs sharing the same dominant haplogroup (Hg I for P14–P6; Hg II/IV for P11–P9). Tando Nceppo and Busogo Beach (P2–P4) were also undifferentiated (Φ_ST_ = −0.094; *p* = 0.948), consistent with shared membership in Hg II and Hg III at both stations.

In contrast, *C. linduensis* displayed no significant among-station differentiation within Lake Lindu (Φ_ST_ = 0.225; *p* = 0.064). All three pairwise comparisons were non-significant before and after Bonferroni correction (Table 2): Nangkating–Uwe Rawa (L1–L2; Φ_ST_ = 0.275; *p* = 0.126), Uwe Rawa–Pulau (L2–L3; Φ_ST_ = 0.222; *p* = 0.208), and Nangkating–Pulau (L1–L3; Φ_ST_ ≈ 0.000; *p* = 1.000). The absence of significant genetic structure among *C. linduensis* stations is consistent with the low haplotype diversity (N*h* = 3) and the dominance of a single haplotype shared across all three stations (Fig. 3A; Table 1).

To evaluate whether haplogroup distributions reflect discrete geographic population units, an additional hierarchical AMOVA was performed partitioning *C. possoensis* sequences into three geographic zones concordant with haplogroup distribution: North [P6 Tentena + P14 Siuri; Hg I], East [P2 Tando Nceppo + P4 Busogo Beach; Hg III], and Southwest [P11 Bancea + P9 Pendolo; Hg II] (Table 2). This analysis attributed 37.0% of total variance to differences among the three geographic zones (Φ_CT_ = 0.370), 5.2% to differences among stations within zones (Φ_SC_ = 0.082), and 57.8% to variation within stations (Φ_ST_ = 0.422, *p* < 0.001). The Φ_CT_ value approached but did not reach conventional significance (*p* = 0.066).

### 3.5. Isolation by distance

The Mantel test revealed no statistically significant relationship between genetic differentiation and geographic distance for either species. For *C. possoensis*, the Mantel statistic was *r* = 0.293 (*p* = 0.144, 9,999 permutations), indicating that the observed population genetic structure was not explained by geographic distances among the six Lake Poso sampling stations. For *C. linduensis*, a high Mantel correlation coefficient (*r* = 0.964) was obtained; however, only three sampling stations were available, yielding an insufficient number of unique permutations for statistical inference (*p* = 0.333).

### 3.6. Demographic history

Tajima’s *D* was negative for both *C. possoensis* (*D* = −0.504, *p* = 0.615) and *C. linduensis* (*D* = −1.022, *p* = 0.307), but neither deviated significantly from zero. Fu’s *F_s_* was likewise negative for both species (*C. possoensis*: *F_s_* = −2.175, *S* = 0.102; *C. linduensis*: *F_s_* = −1.207, *S* = 0.230), indicating a slight excess of haplotypes relative to neutral expectation; however, neither value reached statistical significance (*S* < 0.02). Station-level analyses yielded non-significant Tajima’s *D* at all testable stations within both species (Lake Poso: *D* ranging from −1.062 at P6 to 1.352 at P9; Lake Lindu: *D* = −1.112 at L1 and 0.820 at L2); stations P14 and L3 were excluded due to absence of segregating sites. (Fig. 4C, D)

**Fig. 4.**
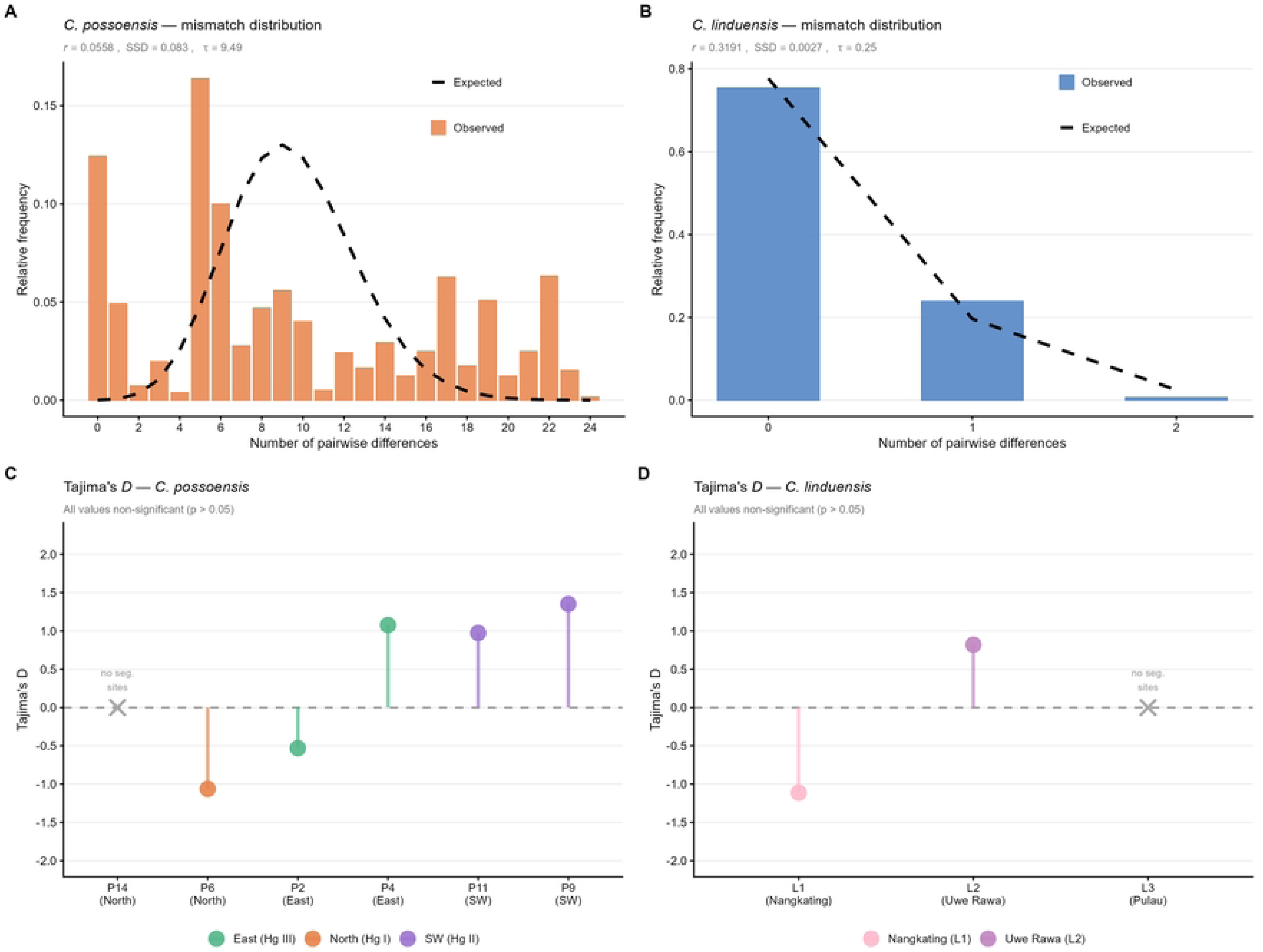
Demographic history inferences for *C. possoensis* (Lake Poso) and *C. linduensis* (Lake Lindu). (A) Mismatch distribution of *C. possoensis* (orange bars) overlaid with the expected distribution under a sudden expansion model (dashed line; Rogers & Harpending, 1992); expansion parameter τ = 9.489, raggedness index *r* = 0.056, SSD = 0.083. (B) Mismatch distribution of *C. linduensis* (blue bars); τ = 0.253, *r* = 0.319, SSD = 0.003. (C) Per-station Tajima’s *D* values for *C. possoensis* across six Lake Poso stations (P2, P4, P6, P9, P11, P14); station P14 was excluded from individual testing due to the absence of segregating sites. (D) Per-station Tajima’s *D* values for *C. linduensis* across Lake Lindu stations (L1, L2, L3); station L3 was excluded due to the absence of segregating sites. Dashed horizontal line indicates neutral expectation (*D* = 0); all values were non-significant (α = 0.05).

The mismatch distribution of *C. possoensis* showed a smooth, unimodal profile (raggedness index *r* = 0.056; SSD = 0.083), with an estimated expansion parameter τ = 9.489. The mismatch distribution of *C. linduensis*, by contrast, was irregular and multimodal (*r* = 0.319; SSD = 0.003), inconsistent with a simple sudden expansion model; the very low τ value (0.253) reflects minimal pairwise divergence among the three observed haplotypes (Fig 4A, B).

## 4. Discussion

### 4.1. Extreme within-species genetic diversity in C. possoensis and the limits of COI-based phylogenetics

The COI haplotype diversity documented in *C. possoensis* (*h* = 0.876; π = 0.0161; 24 haplotypes organized into four geographically structured haplogroups delineated by haplotype network topology) is among the highest recorded for a freshwater bivalve confined to a single lake system, and contrasts sharply with the depauperate variation of *C. linduensis* (*h* = 0.246; π = 0.0004; three haplotypes). Of particular diagnostic relevance, the mean p-distance between geographic zones ranged from 1.41% (Hg I vs. Hg III) to 2.26% (Hg I vs. Hg II), with K2P distances virtually identical throughout (1.44–2.33%; Table 3), confirming that the model correction has negligible effect at this level of divergence (Δ < 0.08% in all comparisons). The maximum between-zone divergence (Hg I vs. Hg II, K2P = 2.33%) approaches the between-species distance between *C. possoensis* and *C. linduensis* (K2P = 2.42%), underscoring that within-lake haplogroup divergence has reached near-interspecific levels. Within-zone K2P values ranged from 0.67% (North/Hg I) to 1.92% (Southwest/Hg II; Table 3). The mean pairwise differences within *C. possoensis* (κ = 9.49 substitutions per 590-bp sequence) thus reflect a level of within-species molecular diversity that, in most freshwater bivalve groups, would span an entire genus.

**Table 3.** Pairwise Φ_ST_ (upper triangle, bold; *** *p* < 0.001, 10,000 permutations) and mean between-zone genetic distances (lower triangle, in %; format: *p*-distance / K2P) for the three geographic haplogroup zones of *C. possoensis*. Diagonal values (grey shading) show within-zone mean *p*-distance / K2P distance. The near-identity of *p*-distance and K2P throughout (Δ < 0.08%) confirms that the model correction has negligible effect at this level of divergence. Zone assignments: North [Hg I: P6 Tentena + P14 Siuri]; Southwest [Hg II: P11 Bancea + P9 Pendolo]; East [Hg III: P2 Tando Nceppo + P4 Busogo Beach].

|  | North (Hg I) | Southwest (Hg II) | East (Hg III) |
| --- | --- | --- | --- |
| North (Hg I) | 0.65% / 0.67% | **0.4423\*\*\*** | **0.4759\*\*\*** |
| Southwest (Hg II) | 2.26% / 2.33% | 1.87% / 1.92% | **0.2759\*\*\*** |
| East (Hg III) | 1.41% / 1.44% | 1.86% / 1.91% | 0.81% / 0.82% |

This extreme divergence directly explains the recurring polyphyletic recovery of *C. possoensis* in COI-based phylogenies. When the four haplogroups within a single morphospecies carry sequences more divergent than typical congeneric species pairs, deep coalescence and incomplete lineage sorting ensure that any gene tree built from COI will reflect lineage history rather than species history (Avise 2000). This problem is not unique to *C. possoensis*: polyphyly attributable to deep intraspecific lineage retention has been extensively documented in freshwater bivalves worldwide, including in the invasive *Corbicula* clades established in European rivers, where multiple genetically divergent lineages of uncertain taxonomic status co-occur and produce reticulate COI topologies that resist resolution by single-locus approaches (Pigneur et al. 2011). A directly comparable case within the Sulawesi ancient lake system is provided by the viviparous freshwater gastropod *Tylomelania*, which underwent a spectacular adaptive radiation in the Malili Lakes, accumulating COI divergences of 5–15% among morphologically similar endemic species (von Rintelen et al. 2004, 2007). In *C. possoensis*, the haplogroup architecture of Lake Poso thus mirrors the incipient radiation process documented in Tylomelania: geographically structured mitochondrial lineages that have accumulated substantial COI divergence within a single enclosed lake system. These lineages appear to be at a stage where morphological differentiation has not yet caught up with molecular divergence.

The same coalescent processes likely account for a second pattern: the placement of *C. anomioides* haplotypes at intermediate positions among the *C. possoensis* haplogroups (Fig. 3C), mirroring its nested recovery in the ML tree (Fig. 2). Such mito-nuclear or morphological discordance between recently diverged, reproductively isolated species is most parsimoniously explained by incomplete lineage sorting of an ancestral mitochondrial polymorphism rather than ongoing gene flow (Maddison 1997, Funk and Omland 2003). Notably, our *C. anomioides* specimens from Dulumai form a well-supported monophyletic cluster, in contrast to the paraphyletic placement of previously published *C. anomioides* sequences; the basis of this discordance — whether geographic, taxonomic, or methodological — warrants dedicated integrative investigation. These observations reinforce that COI alone is insufficient to delimit these endemic lineages.

These findings underscore the fundamental inadequacy of COI alone for species delimitation and phylogenetic reconstruction in Sulawesi *Corbicula*. An integrative approach — combining nuclear loci, detailed morphological characterisation of shell shape and soft tissue anatomy, and ecological data (Bespalaya et al. 2025)— is indispensable for determining whether the haplogroups of *C. possoensis* represent incipient species, deeply structured but interbreeding populations, or ancient lineages maintained as a single morphological taxon by stabilising selection or recent contact. The present study provides the first population-genetic framework for *C. possoensis* and establishes the haplogroup geography that future integrative analyses must resolve. It must be emphasised that the haplogroup designations employed here are purely operational units defined by mitochondrial network topology and geographic concordance; they carry no *a priori* taxonomic weight, and their status as incipient species, structured populations, or retained ancestral lineages cannot be adjudicated on the basis of a single mitochondrial marker alone.

### 4.2. Phylogeographic zonation of C. possoensis within Lake Poso: haplogroup structure as evidence of within-lake historical isolation

Hierarchical AMOVA revealed strong and significant genetic differentiation among *C. possoensis* stations within Lake Poso (Φ∼ST∼ = 0.404, *p* < 0.001). Phylogenetic support for Haplogroup II and III boundaries is strong in the ML tree (SH-aLRT/UFBoot = 84.1/99 and 85/99, respectively), and the haplogroup assignments are further corroborated by the population structure analyses. When stations were grouped according to their dominant haplogroup and geographic position (North: Hg I; East: Hg III; Southwest: Hg II), the haplogroup-based AMOVA attributed 37.0% of total variance to among-group differences (Φ∼CT∼ = 0.370), with only 5.2% among stations within zones (Φ∼SC∼ = 0.082, *p* = 0.074) and 57.8% within stations. The non-significant Φ∼SC∼ demonstrates that the two stations within each geographic zone share a mutually undifferentiated haplotype pool — direct molecular evidence that the three zones represent the primary axes of phylogeographic partitioning within the lake. An analogous pattern of within-lake geographic structuring has been documented in other ancient lake invertebrates of Sulawesi. Among the endemic *Caridina* shrimps of Lake Poso and the Malili system, species from the Malili lakes in particular show pronounced geographic differentiation, with geographically separated populations forming distinct mitochondrial subclades suggesting limited gene flow between shoreline sectors (von Rintelen 2011). Similarly, the epizoic freshwater limpet *Protancylus pileolus* in Lake Poso shows deep phylogeographic structure attributable to drift-based geographic speciation, with genetic data consistent with a geographic speciation mode correlated with present distribution patterns (Albrecht et al. 2020). Strikingly, the basal lineage of *P. pileolus* is restricted to the southern shoreline of Lake Poso — geographically concordant with the Southwest zone of *C. possoensis* (Haplogroup II; Bancea [P11] and Pendolo [P9] stations) — suggesting that the southern basin may represent a phylogeographic refugium shared across ecologically disparate lacustrine taxa. That within-lake geographic barriers produce discrete genetic assemblages in organisms as ecologically diverse as benthic shrimps, epizoic gastropods, and — as shown here — sedentary freshwater bivalves, strongly suggests a shared landscape-level mechanism operating across the Sulawesi ancient lake fauna, rather than taxon-specific dispersal limitation alone.

This geographic concordance between haplogroup distribution and sampling locality is further reflected in the spatial arrangement of the three zones: the North zone (Tentena, P6; Siuri, P14) occupies the northern end of Lake Poso near the Poso River outlet; the East zone (Tando Nceppo, P2; Busogo Beach, P4) occupies the eastern shoreline; and the Southwest zone (Bancea, P11; Pendolo, P9) covers the central-west and southern margins. The absence of a significant isolation-by-distance relationship (Mantel *r* = 0.293, *p* = 0.144; Slatkin, 1993) indicates that this structure is not generated by simple distance-decay in gene flow, but reflects discrete barriers or historical vicariance. Lake Poso is a deep (maximum depth ∼400 m), tectonically formed ancient lake whose steep bathymetric gradients, heterogeneous substrates, and potentially anoxic deep zones along different shoreline sectors could restrict passive movement of a benthic freshwater bivalve with limited vagility (Whitten et al. 2012, Kaban et al. 2023, Damanik et al. 2024). Alternatively, the three-zone structure may preserve the signature of repeated lake-level fluctuations during Pleistocene glacial maxima, when changes in water level may have created habitat barriers between the different shoreline sections of Lake Poso, allowing independent mitochondrial divergence before subsequent lake-level recovery re-established contact without haplotype homogenisation (cf. Voris 2000).

The exceptional position of station P14 (Siuri) deserves specific attention. All ten *C. possoensis* individuals at Siuri shared a single COI haplotype (no segregating sites), despite Siuri being assigned to the same North zone as Tentena (P6), which harbours multiple haplotypes. This within-station monomorphism is consistent either with a recent local founder event — the current Siuri population descending from a very small number of North-zone colonists — or with a past population bottleneck that purged pre-existing variation. Despite its zero internal diversity, P14 showed the highest pairwise Φ_ST_ values against stations from the East and Southwest zones (Table 2), a consequence of its fixed haplotype diverging substantially from the multi-haplotype assemblages elsewhere in the lake. The Siuri population thus contributes a unique and irreplaceable mitochondrial lineage to the overall diversity of *C. possoensis* in Lake Poso, irrespective of its low within-population diversity.

### 4.3. Demographic stasis and panmixia in C. linduensis: a geographically constrained endemic

*C. linduensis* exhibited extremely low COI diversity (*h* = 0.246; π = 0.0004; three haplotypes from 30 individuals across three stations). Within-lake Φ_ST_ (0.225) was non-significant, indicating panmixia across all sampling stations. Nucleotide diversity in *C. linduensis* (π = 0.000429) is approximately 37-fold lower than in *C. possoensis*, and is exceptionally low even by comparison with other threatened freshwater bivalves. For reference, the endangered freshwater pearl mussel *Margaritifera margaritifera* — one of the most extensively studied imperilled freshwater bivalves globally — displays nucleotide diversity values of up to π = 0.004 across its fragmented European populations (Zanatta et al. 2018), an order of magnitude higher than *C. linduensis*. The only comparable COI diversity levels among freshwater bivalves are found in invasive androgenetic *Corbicula* lineages, which are effectively clonal and thus represent a fundamentally different biological context (Peñarrubia et al. 2017, Vastrade et al. 2022, Ludwig et al. 2024). Among sexually reproducing freshwater bivalves with intact natural populations, the COI diversity of *C. linduensis* thus represents one of the lowest values documented to date, just equal to or almost similar to some freshwater bivalves that have experienced severe demographic bottlenecks or extreme habitat isolation. For instance, the sexually reproducing Wabash pigtoe, *Fusconaia flava*, shows a heavily restricted COI nucleotide diversity (π = 0.00078) in the Green River system (Olivera-Hyde et al. 2023). Similarly, the endangered scaleshell mussel, *Leptodea leptodon*, exhibits substantially lower mitochondrial genetic diversity compared to its more common congener *L. fragilis* due to high habitat specificity and historical population decline (Chong and Roe 2018). Furthermore, within the genus *Corbicula* itself, the sexually reproducing brackish-water species *C. japonica* has demonstrated restricted genetic diversity dictated by its specific reproductive structure and geographical isolation across different lake systems (Mito et al. 2014, Yamada et al. 2014). These comparisons highlight that while the extreme genetic homogeneity of *C. linduensis* is rare for a sexually reproducing bivalve, it aligns with the profound genomic erosion typically observed in highly isolated populations persisting at the lowest limits of genetic variation.

Two non-exclusive hypotheses account for this pattern. Under a ‘young colonist’ scenario, *C. linduensis* colonised Lake Lindu relatively recently from an ancestral Sulawesi Corbicula stock — perhaps via passive dispersal by migratory waterbirds, a documented long-range dispersal vector for freshwater bivalves (Bilton et al. 2001, Barboza et al. 2022) — and insufficient time has elapsed for substantial haplotype diversification. Under a ‘small effective population’ scenario, the lineage has resided in Lake Lindu for a longer period, but its small effective size, imposed by the lake’s restricted area (∼30 km²) and geographical isolation enforced by the surrounding Central Sulawesi Highlands (Sutapa et al. 2018), has progressively eroded mitochondrial diversity through genetic drift. The mismatch distribution supports the latter interpretation: the irregular, multimodal profile (raggedness index *r* = 0.319; Harpending 1994) and very low τ (0.253) are inconsistent with sudden expansion (Rogers and Harpending 1992) and instead reflect a stable, small-*N*_e_ population in which most pairwise comparisons yield zero or one substitution. Under either hypothesis, *C. linduensis* has not undergone the long-term within-lake diversification that shaped the haplogroup architecture of *C. possoensis* in Lake Poso.

Panmixia within Lake Lindu is consistent with the lake’s modest dimensions: the maximum inter-station distance is far smaller than in Lake Poso, and passive dispersal — whether by currents, aquatic vertebrates, or waterbirds — would readily homogenise allele frequencies across the lake basin (Bilton et al. 2001). The high Mantel correlation (*r* = 0.964) is statistically uninterpretable with only three stations (six unique permutations; *p* = 0.333; Slatkin 1993) but suggests that a denser sampling scheme might reveal subtle distance-structured variation. The near-absence of genetic structure in *C. linduensis* stands in stark contrast to the pattern in *C. possoensis*, reinforcing the conclusion that lake size and isolation history — not just taxonomic affinity — are the primary determinants of within-lake phylogeographic architecture in these Sulawesi endemics.

### 4.4. Deep vicariance between Lake Poso and Lake Lindu: ancient divergence confirmed by hierarchical AMOVA

The three-level AMOVA of the combined dataset revealed that 60.7% of total COI variance is attributable to between-lake differences (Φ_CT_ = 0.607, *p* < 0.001), dwarfing the within-lake Φ_ST_ for either species (0.404 for *C. possoensis*; 0.225 ns for *C. linduensis*). Lake Poso and Lake Lindu have no hydrological connection: the Poso River drains northward to Tomini Bay while Lake Lindu’s catchment empties separately to the east/south-east (Whitten et al. 2012). Under prolonged geographic isolation within the tectonically active Central Sulawesi landscape, independent mitochondrial evolution in the two respective *Corbicula* lineages is expected to generate exactly the deep divergence documented here. The profound genetic discontinuity between the two species (ΦCT = 0.607) mirrors the deep between-lake mitochondrial divergences reported for endemic gastropods partitioned among the Malili Lakes of Sulawesi (Von Rintelen et al. 2004, Glaubrecht and Von Rintelen 2008), where ancient geographic barriers have similarly restricted gene flow and produced deep interspecific-level divergence from shared ancestors. Congruent patterns of deep vicariance and pronounced genetic structure across lake boundaries have also been extensively documented in other Sulawesian lacustrine taxa, including atyid shrimps (Von Rintelen et al. 2010a, von Rintelen 2011) and pelagic copepods (Vaillant et al. 2013).

The between-lake divergence provides strong molecular corroboration for recognising *C. possoensis* and *C. linduensis* as distinct species, complementing their morphological differentiation. The clear two-tier hierarchical structure — deep interspecific divergence (ΦCT = 0.607) superimposed on within-species among-station differentiation (ΦSC = 0.423) — is consistent with two lineages that have undergone prolonged independent diversification within ancient, hydrologically isolated lake environments. Importantly, the available phylogenetic data do not support a direct sister-group relationship between *C. possoensis* and *C. linduensis*; the ML tree places *C. linduensis* in closer affinity to the Malili lake species (*C. loehensis*, *C. matannensis*) than to *C. possoensis*, suggesting that the two Lake Poso and Lake Lindu lineages were derived from separate ancestral stocks rather than from a single vicariance event separating the two lakes. The deep between-lake divergence is therefore better interpreted as the outcome of long-term allopatric isolation acting independently on each lineage within its respective lake, rather than as evidence of direct allopatric speciation between *C. possoensis* and *C. linduensis* sensu stricto. This model of independent lacustrine diversification under geographic isolation is consistent with the ancient-lake vicariance framework previously reported for the Malili Lake system (Von Rintelen et al. 2010b), and appears to characterise the broader pattern of endemic freshwater diversification across the ancient lake systems of Central Sulawesi.

### 4.5. Contrasting demographic histories and the confounding effect of haplogroup structure on neutrality tests

The mismatch distribution of *C. possoensis* showed a smooth, unimodal profile (raggedness index *r* = 0.056; Harpending 1994) superficially consistent with the Rogers and Harpending 1992 model of past sudden population expansion (τ = 9.489). However, interpreting this profile as a direct demographic expansion signal requires caution in the context of the species’ complex haplogroup structure. When sequences from multiple divergent haplogroups are pooled, the observed pairwise differences comprise two superimposed distributions: a low-κ component from within-haplogroup comparisons and a high-κ component from between-haplogroup comparisons. The between-haplogroup component can dominate and shift the distribution toward a broad, apparently unimodal shape — mimicking an expansion signal even in the absence of genuine demographic growth. This confound is well recognised in phylogeographic studies where multiple divergent lineages are sampled together (cf. Excoffier 2004). Future analyses employing haplogroup-specific mismatch distributions, extended Bayesian skyline plots per geographic zone, and full-coalescent inference would disentangle genuine demographic expansion from a lineage-pooling artifact.

In contrast, the demographic inference for *C. linduensis* is unambiguous. With only three haplotypes differing by at most one or two substitutions, the irregular mismatch (*r* = 0.319) and minimal τ (0.253) are consistent with long-term demographic stasis in a small, stable population — a pattern that mirrors the genetic signatures reported for other highly restricted freshwater endemics from small island lakes (Dawson and Hamner 2008). Tajima’s *D* (−1.022, *p* = 0.307; Tajima, 1989) and Fu’s *F_s_* (−1.207, *S* = 0.230; Fu, 1997) are non-significant but directionally consistent with a small stable population: the near-absence of segregating sites severely limits the power of these tests, and the non-significance should not be interpreted as evidence against any particular demographic scenario.

More broadly, the non-significance of Tajima’s D and Fu’s Fs at the species level for *C. possoensis* (Tajima 1989, Fu 1997) reflects the well-documented sensitivity problem of single-population neutrality tests when applied to population-structured samples. Both statistics assume a single panmictic population; when applied to samples drawn from multiple differentiated haplogroups, among-lineage variance inflates the test statistic variance and substantially reduces power (Ptak and Przeworski 2002). The negative direction of both statistics — suggesting excess rare variants or excess haplotypes relative to neutral expectation — is consistent with an expansion or purifying selection signal, a demographic pattern comparably inferred from negative neutrality indices in other *Corbicula* taxa, such as the estuarine *C. japonica* (Mito et al. 2014) and the widespread *C. fluminea* (Zhu et al. 2018). Therefore, the failure to reach significance does not constitute evidence against demographic change. Instead, it reflects the structural limits of applying single-population models to a deeply subdivided dataset, a confounding factor frequently encountered when interpreting the demographic history of structured lacustrine populations in Wallacea (Vaillant et al. 2013).

### 4.6. Conservation implications: distinct management units and the vulnerability of endemic lake bivalves

The phylogeographic zonation in *C. possoensis* has direct implications for conservation management. The three geographic zones — North, East, and Southwest — are characterised by distinct dominant haplogroups with significant pairwise ΦST (Table 2) and p-distances of 1.41–2.26% (K2P: 1.44–2.33%; Table 3) between zones. Rather than definitively classifying them as Evolutionarily Significant Units (ESUs) based solely on a single mitochondrial locus, it is more appropriate and conservative to provisionally treat each zone as a distinct Management Unit (MU; Moritz, 1994). For conservation management, these three MUs should be managed as discrete entities: translocation of individuals between zones risks introducing non-local haplotypes into resident populations and disrupting locally adapted gene complexes (Frankham et al. 2002). This caution is especially pertinent for station P14 (Siuri), whose monomorphic population would be permanently altered by any introgression of foreign haplotypes.

For *C. linduensis*, the priority is not preserving genetic structure — which is absent— but preventing further genetic erosion in an already depauperate species. With only three mitochondrial haplotypes across the entire range and one station (L3, Pulau) already monomorphic, the current gene pool of *C. linduensis* is extremely vulnerable to stochastic loss. Lake Lindu’s small area (∼30 km²) means that a single lake-wide disturbance event — a toxic algal bloom, major sedimentation pulse, or invasion by the globally problematic *Corbicula fluminea* (Sousa et al. 2014) — could simultaneously threaten the entire species. We recommend that *C. linduensis* be formally assessed under IUCN Red List criteria, with particular attention to the Restricted Range and Small Population sub-criteria, and that genetic diversity metrics (haplotype richness, nucleotide diversity) be incorporated into long-term monitoring alongside abundance estimates.

Furthermore, while the present COI baseline is foundational, multi-marker and genomic reassessments (e.g., genome-wide SNPs or whole-genome resequencing) are urgently needed for both *C. possoensis* and *C. linduensis*, as well as other endemic Sulawesian *Corbicula*. Transitioning from single-locus to genomic data is critical to accurately delineate true ESUs, quantify genome-wide inbreeding and effective population sizes (Ne), and assess the adaptive potential of these highly isolated populations under changing environmental conditions (Kardos et al. 2021, Perea et al. 2022, Faust et al. 2025).

Lake Poso is a well-established ancient lake (von Rintelen et al. 2004, Vaillant et al. 2013), whereas Lake Lindu is currently considered a putative ancient lake whose precise geological antiquity requires further corroboration (Annawaty et al. 2016). Nevertheless, both of these tectonic lakes harbour exceptionally high aquatic endemism. This includes endemic fish radiations such as the remarkable diversity of ricefishes (Parenti 2008, Sutra et al. 2019, Möhring et al. 2025), endemic gastropods (von Rintelen et al. 2004, 2014, Glaubrecht and von Rintelen 2008), and the *Corbicula* clams studied here. The genetic baselines established in this study — the first population-level analyses of *C. possoensis* and *C. linduensis* — provide essential reference data for future conservation genetics assessments and for monitoring the impacts of ongoing environmental pressures. These pressures include the spread of alien invasive species, introduction of exotic fish, increasing aquaculture, and land-use change in the surrounding watershed (Herder et al. 2022, Rahmawati et al. 2025). Preserving the full extent of mitochondrial diversity documented here — across all three geographic zones in Lake Poso and the entirety of Lake Lindu — requires protection of multiple shoreline sites and the maintenance of hydrological integrity in both lake ecosystems.

## 5. Conclusion

This study establishes a foundational population-genetic framework for the endemic *Corbicula* of Sulawesi. Our findings reveal two fundamentally contrasting evolutionary trajectories shaped by the landscape of Central Sulawesi. *Corbicula possoensis* in Lake Poso is characterised by exceptionally high genetic diversity and pronounced micro-geographic structuring, with divergence between its phylogeographic zones approaching the level separating it from *C. linduensis* — a pattern echoing the deep intra-lacustrine differentiation documented in other Sulawesian endemic fauna. In stark contrast, *C. linduensis* exhibits profound genetic depauperation and demographic stasis, underscoring the vulnerability of endemic species restricted to smaller, putatively younger or more isolated lake basins such as Lake Lindu. Notably, the absence of a direct sister-group relationship between the two species indicates that their divergence reflects independent diversification within separate lake systems rather than a single vicariance event, consistent with the broader pattern of endemic radiation across the ancient lakes of Central Sulawesi.

From a conservation perspective, the distinct spatial haplogroups of *C. possoensis* should be provisionally recognised as discrete Management Units (MUs) to prevent the disruption of local genetic integrity through human-mediated translocation. For *C. linduensis*, immediate conservation monitoring is imperative to protect its homogeneous and fragile gene pool from stochastic events and the looming threat of invasive species. Finally, while this COI-based assessment reveals deep historical isolation, we strongly advocate future multi-marker and whole-genome reassessment. Moving from single-locus to genome-wide data is an essential next step to definitively resolve species boundaries — including the phylogenetic nestedness of *C. anomioides* within *C. possoensis* — assess genome-wide inbreeding, and refine conservation strategies for the irreplaceable freshwater biodiversity of the Wallacea region.

## Acknowledgement

This project was financially supported by a research grant from the Nagao Natural Environment Foundation (NEF) awarded for the period of 2025–2026 and partly supported by the Research and Innovation for Advanced Indonesia (RIIM BRIN-LPDP) for the publication. We thank the National Research and Innovation Agency (BRIN) of the Republic of Indonesia for issuing the necessary research and ethical permits, as well as the *Tua Adat* of the *To‘oma To‘Lindu* indigenous community for their permission and guidance during the fieldwork. Finally, we express our deepest gratitude to all colleagues, field assistants, local fishers, and community members who contributed to the fieldwork and logistical support. Although they cannot be mentioned individually, their assistance was invaluable to the successful completion of this study.

